# Acid-Base Homeostasis and Implications to the Phenotypic Behaviors of Cancer

**DOI:** 10.1101/2022.03.04.482927

**Authors:** Yi Zhou, Wennan Chang, Xiaoyu Lu, Jin Wang, Chi Zhang, Ying Xu

## Abstract

Acid-base homeostasis is a fundamental property of living cells and its persistent disruption in human cells can lead to a wide range of diseases. We have conducted computational modeling and analysis of transcriptomic data of 4750 human tissue samples of nine cancer types in the TCGA database. Built on our previous study, we have quantitatively estimated the (average) production rate of OH^−^ by cytosolic Fenton reactions, which continuously disrupt the intracellular pH homeostasis. Our predictions indicate that all or a subset of 43 reprogrammed metabolisms (RMs) are induced to produce net protons (H^+^) at comparable rates of Fenton reactions to keep the intracellular pH stable. We have then discovered that a number of well-known phenotypes of cancers, including increased growth rate, metastasis rate and local immune cell composition, can be naturally explained in terms of the Fenton reaction level and the induced RMs. This study strongly suggests the possibility to have a unified framework for studies of cancer-inducing stressors, adaptive metabolic reprogramming, and cancerous behaviors. In addition, strong evidence is provided to demonstrate that a popular view of that Na^+^/H^+^ exchangers, along with lactic acid exporters and carbonic anhydrases are responsible for the intracellular alkalization and extracellular acidification in cancer may not be justified.

## Introduction

Acid-base homeostasis is a most fundamental property that all living cells must maintain as the pH sets the stage for performing accurately all the biochemistry needed in support of the livelihood and the functionalities of the cells. pH is an essential part of probably all cellular processes of a living organism, which ensures the correct folding of proteins, biomolecular binding and interactions with the right affinity and specificity, conducting reactions at the desired rates in enzymatic pathways, among other biological functions. For normal human tissue cells, the intracellular (or cytosolic) pH, pH_*i*_, is generally neutral or slightly acidic at the range of 6.8–7.1 while the extracellular pH, pH_*e*_, is slightly basic at ~7.2 with pH_*i*_ < pH_*e*_ (1). Individual cellular compartments may have distinct pH levels required for executing their functions, with lysosomes having the most acidic pH at ~4.0-5.0 and mitochondria having the highest one at ~8.0 (2). Because of the vital importance of the pH, all living cells have designated systems to keep the stability of their pH_*i*_ and pH_*e*_.

In human cells, a HPO_4_^2−^/H_2_PO_4_^−^-based and a HCO_3_^−^/H_2_CO_3_-based buffering system are used to maintain the stability of pH_*i*_ and pH_*e*_, respectively, plus a suite of proton importers and exporters such as V-ATPase, Na^+^/H^+^ exchanger and Na^+^/HCO_3_^−^ symporter. Under physiological conditions, certain metabolisms may produce large quantities of protons such as *de novo* nucleotide biosynthesis (3) and the Warburg effect (4) while some other metabolisms may consume protons such as the conversion of NADH to NAD^+^ (5). Excess protons or hydroxides (or equivalents) produced dynamically by such acidifying or alkalizing processes are generally absorbed by the pH buffer and/or counterbalanced by proton transporters to keep the stability of the pH.

Persistent pathological conditions such as chronic inflammation are known to disrupt the stability of the local pH; and extracellular acidosis has been widely reported in (6–8). For example, cells in Alzheimer’s disease tissues are known to be under both extracellular and intracellular acidosis (9, 10). Similar observations have been reported about other neurodegenerative diseases (11). Diabetes is another example where the diseased tissue cells have been reported to have more acidic extracellular pH than the matching healthy tissues (12, 13). Knowing the vital importance of the cellular pH stability, one could imagine the profound impacts of such changes on the whole cellular biochemistry. This is the reason that persistently altered pH has been suggested as a fundamental cause to a wide range of the altered metabolisms, hence considerable cellular behaviors, in neurodegenerative diseases, diabetes, and cancer.

It is noteworthy that for pathological conditions giving rise to persistent overproduction of protons or hydroxides, the pH buffer, along with the proton transporters, has only a limited power in maintaining the pH stability. The reason is twofold: (1) each pH buffer has a fixed capacity, which could absorb only limited protons (or hydroxides) (14, 15) persistently generated under pathological conditions; and (2) proton transporters generally do not work in a sustained manner as such transporters fall into two types, *electroneutral* co/antiporters and *electrogenic* transporters. For electroneutral co/antiporters, protons are released from or loaded into cells at the expense of extruding or absorbing another ion, e.g., Na^+^ or Cl^−^. Hence their persistent utilization will disrupt the homeostasis of the other ion, making them not a sustainable solution. Similar can be said about a H^+^ importer or exporter (or equivalent), since it is electrogenic and its persistent utilization will violate the *electroneutrality* of the host cells, another fundamental property that cells must keep to remain viable (16).

Cancer is an intriguing case in terms of the altered pH_*i*_ and pH_*e*_ as it has been well established that the pH_*i*_ of cancer tissue cells becomes basic, at ~7.4 or 7.5 while their pH_*e*_ becomes acidic ranging from 6.4 to 6.8 (1), hence a reversal of pH_*i*_ < pH_*e*_ compared to normal tissue cells. Multiple proposals have been made regarding the possible causes for the reversal of the pH_*i*_ and pH_*e*_ levels. These include (i) upregulated proton exporters such as V-ATPase, Na^+^/H^+^ exchangers, and lactic acid exporters in cancer (17–19); (ii) increased utilization of carbonic anhydrases that convert extracellular CO_2_ released by cancer cells to HCO_3_^−^ and H^+^ (20, 21); (iii) hypoxia due to poor blood supply and “respiratory bursts” by innate immune cells (22). These proposals have addressed the possible reasons for the extracellular acidosis in cancer tissues, which is probably needed by the local immune cells (6) but did not actually answer the question: *what has made the intracellular pH alkaline* as we will demonstrate in this report.

V-ATPase is generally employed in the membrane of intracellular compartments and used to acidify compartments like endosome or lysosome (23, 24). It is used in plasma membrane for acidification of the extracellular space only in specialized cells such as osteoclasts and kidney cells. However, there have been no experimental data supporting the proposal that V-ATPase is localized in plasma membrane of cancer tissue cells, to the best of our knowledge, except by studies reporting that the proton pump is localized to the plasma membrane of certain metastasizing cancer cell lines (25, 26). Na^+^/H^+^exchangers are an interesting case, which are driven by both the gradients of Na^+^ and H^+^ with Na^+^-in being with the gradient and H^+^-out against the gradient when reversing pH_*i*_ and pH_*e*_. We will demonstrate here that the potential generated by Na^+^-in is not sufficient to drive H^+^-out in cancer tissues. Lactic acid (CH_3_CH(OH)CO_2_^−^ + H^+^) exporters like MCT1 are used by possibly all cancers in a sustained manner, as long as the acid is continuously produced by cancer cells (27). We will demonstrate that for cells relying on the Warburg effect for ATP production, lactic acid exporters do not contribute to intracellular alkalization.

Overall, the existing proposals did not adequately answer the question about the observed reversal of the pH_*i*_ and pH_*e*_ levels. Hence further studies are needed. We have previously proposed a fundamentally different reason for the considerable alkalization of the intracellular pH in cancer tissue cells for most, possibly all cancer types (28).

Chronic inflammation is known to be causally linked to cancer onset and development (29), giving rise to increased local concentrations of H_2_O_2_. Once the H_2_O_2_ concentrations reach beyond a certain level, local red blood cells may become senescent due to the oxidation of their plasma membranes and their lack of a membrane repair mechanism (30), leading to their engulfment by macrophages (30) and local accumulation of irons released by macrophages after engulfment (31). Under the condition of immune responses, local epithelial cells will sequester the free irons (32), leading to overload of intracellular irons. It has been widely reported that multiple chronic inflammatory diseases (33) and many, possibly all cancer tissues have iron overload (34). It is noteworthy that when both the H_2_O_2_ and Fe^2+^ levels are sufficiently high, Fenton reaction: Fe^2+^ + H_2_O_2_ → Fe^3+^ + HO^•^ + OH^−^, an inorganic reaction without involving any enzyme, will happen (35), regardless being cancer or non-cancerous tissues. Fenton reactions have been widely observed in cancer tissues (36–38), and their levels are generally considerably higher in cancer vs. related non-cancerous disease tissues as shown in Figure S1. Our previous work has discovered that all cancer tissue cells have Fenton reactions in their cytosol, mitochondria, extracellular matrix, and cell surface, respectively, and the reactions will continue if there are reducing molecules nearby that can convert Fe^3+^ back to Fe^2+^ such as superoxide (O_2_^•−^), NADH or Vc (28). In addition, all cancers use O_2_^•−^, generated by local immune cells and mitochondria of the cancer cells, as the main reducing molecules, which drives the cytosolic Fenton reactions continuously in the following form: O_2_^•−^ + H_2_O_2_ → HO^•^ + OH^−^ + O_2_, referred to as the *Haber-Weiss reaction* with Fe^2+^ serving as a catalyst (39, 40). Furthermore, the rates of the cytosolic Fenton reactions in cancer can quickly saturate the intracellular pH buffer, hence driving the cytosolic pH up if the persistently produced OH^−^ is not neutralized. It is noteworthy to emphasize that persistent intracellular Fenton reaction is the result of chronic inflammation coupled with local iron overload, without involving any enzymes. One reliable way for computationally estimating the level of (cytosolic) Fenton reaction is through checking the level of 20S proteasome genes. The 20S proteasome is solely responsible for degradating protein aggregates formed due to interaction between hydroxyl radical (HO^•^) and proteins, where HO^•^ can only be produced intracellularly by Fenton reaction (28).

Now the question is: how do such Fenton reaction-affected cells keep their pH_*i*_ within a viable range? We have previously proposed a model regarding how cancer tissue cells reprogram numerous metabolisms (RMs) to produce protons together at rates comparable to those of the cytosolic Fenton reactions, hence keeping their intracellular pH stable. The model is strongly supported by the observation that each of these RMs is found to produce more protons or consume fewer protons than its original metabolism (41). The key RMs include (a) *de novo* biosynthesis of nucleotides; (b) the Warburg effect for ATP production; (c) simultaneous biosynthesis and degradation of triglyceride; and (d) overproduction and deployment of sialic acids and gangliosides. Here, we present a computational modeling analysis to provide further evidence that the RMs in each cancer tissue are indeed induced to produce protons collectively at a rate comparable to the average rate of the cytosolic Fenton reaction.

We also demonstrate that phenotypes known to be associated with specific cancer (sub)types can be naturally explained in terms of the induced RMs.

## Results

We have conducted a modeling analysis to estimate quantitatively the level of cytosolic Fenton reaction in each cancer tissue and a regression analysis to predict which RMs are induced to produce protons to keep the intracellular pH stable in the sample, followed by an association analysis between the known phenotypes or specific (sub)types and selected RMs in the relevant tissue samples. Overall, we applied our analyses to 4750 cancer samples, along with 503 matching control samples, in nine cancer types (11 subtypes), all based on transcriptomic data from the TCGA database and single cell RNA-seq data (scRNA-seq) of head and neck cancer and melanoma to validate our results.

### pH reversal by transporters?

Multiple proposals have been made regarding the possible causes of intracellular alkalization and extracellular acidification in cancer. One popular model is that Na^+^/H^+^ exchangers, particularly NHE1 (SLC9A1), are the main reason for the reversal of pH_*i*_ and pH_*e*_ in cancer tissues, along with monocarboxylate-H^+^ efflux cotransporters MCT1 and MCT4 (SLC16A1 and SLC16A3) and carbonic anhydrase for CO2 hydration (1). Here we demonstrate that this possibility is low.

We have noted that among the nine cancer types under study, SLC9A1 is upregulated in only three (BRCA, HNSC, THCA) and downregulated in six types, as detailed in Table S1, indicating that SLC9A1 may not play a key role in most of the cancer types.

While ATP is known to be involved in the activation of SLC9A1, ATP is not involved in driving the transport (42). Hence the transporter is driven solely by gradients. Note that reversal of pH_*i*_ and pH_*e*_ requires the transporter to move the intracellular H^+^ against the gradient out of the cell, indicating that the act must be driven by the gradient between the extracellular and intracellular Na^+^ concentrations. It is known that the normal intracellular sodium concentration ranges from 10 to 15 mmol/L, hence 12 mmol/L being used here, and the extracellular one is 140 mmol/L (43). The total sodium concentration (TSL) in cancer (TSL_*C*_) is generally 2–3 fold of the matching normal one (TSL_*N*_) (44, 45). The ratio between the extracellular and intracellular volume in a unit tissue is 20:80 (46). Our goal is to estimate the intracellular sodium concentration (ISL) in cancer (ISL_*C*_), assuming the extracellular sodium concentration (= blood sodium concentration) remains roughly stable. Hence, we have

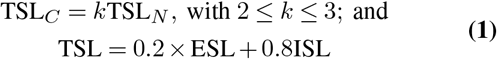

By plugging the relevant numbers, we have

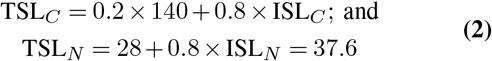

For *k* = 2, we have TSL_*C*_ = 28 + 0.8 × ISL_*C*_ = 2TSL_*N*_. = 75.2. So ISL_*C*_ = 47.2/0.8 = 59. For *k* = 3, we have ISL_*C*_ = 106. We conclude that the intracellular sodium concentration of a cancer tissue is on average 4.91 to 8.83-fold of that of the matching normal tissue. Therefore, the intracellular sodium concentration in cancer should range from 4.91 × 12 ≈ 59 mmol/L to 106 mmol/L. Hence, the free energy gained for moving Na^+^ into the cells from the extracellular space for cancer cells can be calculated as

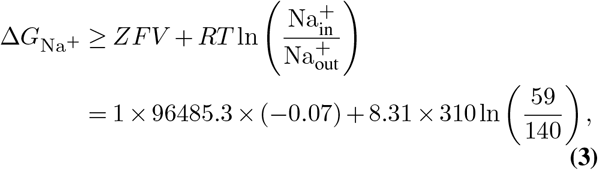

where *Z* =1; *F* = 96485.3 is the Farady constant; and *V* is the transmembrane potential (−0.07 meV inside membrane and 0 outside the membrane); and *R* is the gas constant and *T* is the temperature (room temperature). Similarly, the free energy needed for moving an intracellular H^+^ out of a cancer cell is, using pH_*e*_ = 6.6 and pH_*i*_ = 7.4:

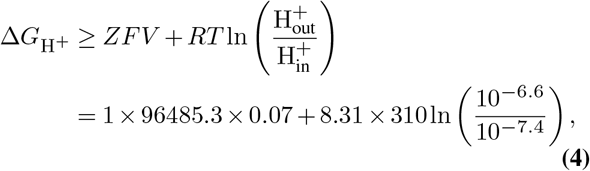

Therefore, the total free energy for Na^+^-in and H^+^-out is Δ*G*_Na+_ + Δ*G*_H+_. Note that the first terms in the two free energies cancel each other, and the total free energy is:

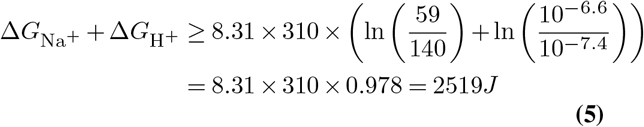

The positive free energy indicates that the energy generated by Na^+^ import is not sufficient to change the pH_*e*_ to 6.6 and the pH_*i*_ to 7.4, actually not even to pH_*e*_ = 6.8 and pH_*i*_ = 7.2 by SLC9A1. This calculation result is also experimentally supported (42). It is noteworthy that we are using the lower bound for the intracellular sodium concentration in cancer. Hence a higher level of such concentration will make it more unlikely for the sodium gradient to drive the reversal of pH_*e*_ and pH_*i*_. Furthermore, the blood sodium levels in cancer patients are generally reduced, a widely known fact (47), suggesting that the actual Δ*G*_Na+_ + Δ*G*_H+_ is higher than the number given in Eq. (5).

Interestingly, the reversal of pH_*e*_ and pH_*i*_ is potentially achievable by SLC9A1 in normal tissue cells, where the ratio between intracellular and extracellular Na^+^ is 12:140 with

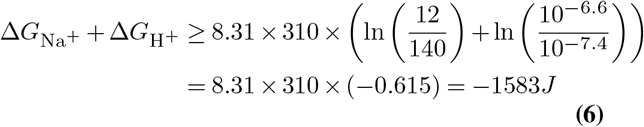

suggesting the possibility of reversing pH_*e*_ and pH_*i*_ by SLC9A1.

The conclusion here is that SLC9A1 could not accomplish the observed reversal of pH_*e*_ and pH*¿* because the Na^+^-in potential is considerably reduced in cancer due to the decreased ratio between intracellular and extracellular Na^+^ concentrations.

Lactic acid exporters SLC16A1 and SLC16A3 have also been suggested to play a role in intracellular alkalization in cancer. We can see from the following that this also is not true, when coupled with the Warburg effect. Note that ATP production by fermenting glucose is pH neutral as given below (48):

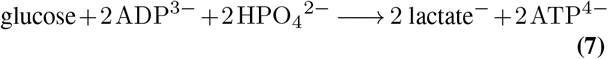

which generates only a lactate but not lactic acid (lactate + H^+^) per ATP produced. Now, the question is where the H^+^comes from when SLC16A1/3 releases a lactic acid. Note that ATP hydrolysis (or consumption) produces one net H^+^ regardless of how the ATP is produced:

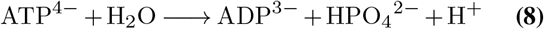

Hence the Warburg effect coupled with ATP hydrolysis produces one net H^+^, which is co-released with the lactate. As a side note, ATP production by respiration consumes one H^+^ for each ATP produced:

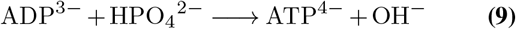

Hence ATP production by respiration coupled with ATP hydrolysis is pH neutral. This is a fundamental difference between the two ATP production pathways.

We conclude: while SLC16A1/3 contributes to the extracellular acidification, it does not contribute to the intracellular alkalization. Our previous work has provided strong evidence that cancer cells release the lactic acids mainly for modulating immune responses rather than pH homeostasis (49).

Carbonic anhydrases, particularly CA4 and CA7, have been suggested to play important roles in extracellular acidification in cancer. As we can see from Table S2, *CA4* is either downregulated or expressed at very low level across all cancer types except for STAD, where the expression increases a little but remains at a very low level. Similarly, *CA7* is down-regulated or expressed at very low level except for THCA. These indicate that the two genes do not play much role in extracellular acidification in all the nine cancer types.

### Estimation of hydroxide production rates by cytosolic Fenton reactions

Our goal here is to construct a reliable metabolic network leading to cytosolic Fenton reaction and to estimate accurately the rate of the hydroxide (OH^−^) production by the Fenton reaction based on transcriptomic data of the available cancer tissues.

To model the rate of the persistent cytosolic Fenton reaction: O_2_^•−^ + H_2_O_2_ → HO^•^ + OH^−^ + O_2_ with Fe^2+^ as the catalyst, we need to estimate the concentration of each of the three reactants: H_2_O_2_, O_2_^−^, and Fe^2+^ and how each product is related to the reactants. **Figure 1A** depicts our constructed map of iron metabolic reactions in a human cell, which consists of three sources to the cytosolic Fe^2+^ pool, namely ferrous ion import, ferric ion import, and reduction, and heme import and reduction; four sinks for the cytosolic Fe^2+^, namely mitochondrial Fe–S cluster and heme synthesis, ferrous ion export, and Fenton reaction; and sources and sinks of the O_2_^−^ and H_2_O_2_, totaling eight. The fifteen metabolic branches were each considered as a *metabolic module*, each containing one to a few dozen of metabolic genes, whose expression levels were utilized to estimate their metabolic flux. Detailed information about gene names and rationale is given in Table S3.

**Fig. 1.**
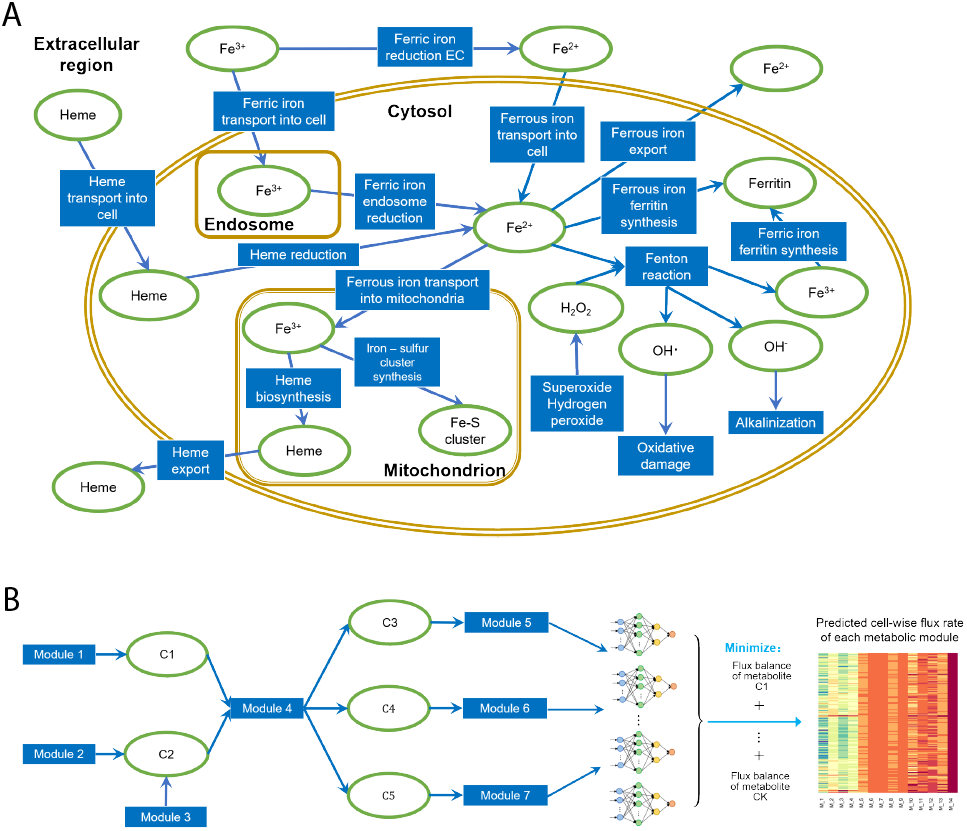
Estimating Fenton reaction rates. **A.** A predicted map for iron metabolism relevant to cytosolic Fenton reaction in human cell. Reactions and metabolites are represented by blue rectangles and green ovals, respectively. **B.** Computational model of scFEA. Each metabolic reaction (or module) is modeled as a neural network of genes involved in the module. The parameters of the neural network were derived by minimizing the total flux imbalance of intermediate metabolites, an indicator for the quality of a predicted model.

We have recently developed a graph neural network-based method for predicting sample-specific metabolic rate, named *scFEA* (single-cell flux estimation analysis) (50). Specifically, scFEA models metabolic fluxes in each tissue based on gene-expression data of a large number of tissue samples, under a few simple and reasonable assumptions. They are (1) the total influx of each metabolite is approximately the same as its total outflux; and (2) changes in the outflux of each metabolite can be modeled as a (non-linear) function of changes in the expression levels of genes involved in producing the metabolite. Note that assumption (1) is generally true unless some major in-/out-flux for a metabolite is not considered. Assumption (2) is a combination of two simpler assumptions: (i) the concentration of an enzyme is an (inversible) function of the concentrations of its reactants; and (ii) this concentration is also a (non-linear) function of the expression level of its encoding gene, with both functions being invariant across different samples of different cancer types. Both assumptions (i) and (ii) are supported by published studies (51–53). Intuitively, one can think this model as an integrated Michaelis-Menten model, whose parameters are implicitly estimated using the large number of available gene-expression data.

Figure 1B outlines the workflow with further details given in Materials and Methods. Specifically, based on the two assumptions, scFEA models the metabolic flux of each module by a three layer fully connected neural network of genes involved in the module, which minimizes the total imbalance of the intermediate substrates across all tissue samples. For a network with *X* modules, there are 12*X* × (# genes) unknowns to be estimated with (# genes) being the number of genes encoded in each reaction-representing module, which is generally a small integer; and there are *K* × *N* constraints, where *K* and *N* are the numbers of intermediate substrates and samples, respectively. Hence a network like the one in Figure 1A, a transcriptomic dataset of more than 2000 samples such as the TCGA pan-cancer (and two selected scRNA-seq) data, should enable reliable estimation of the unknowns. We have validated the scFEA algorithm on human global metabolic map and central metabolic pathways by using two sets of matched scRNA-seq and tissue metabolomics data (50). Here we further validated scFEA on the curated iron ion metabolic modules by applying the method on our recently collected scRNA-seq data of 168 patient-derived pancreatic cancer cell lines Pa03c under four conditions: normoxia (N), hypoxia (H), normoxia and knock-down of APEX1 (N-APEX1-KD), and hypoxia and knock-down of APEX1 (H-APEX1-KD). APEX1 plays a central role in the cellular response to oxidative stress (54). Our recent studies identified that knockdown of APEX1 results in increased oxidative stress and cell death in Pa03c cells (55). scFEA predicts the level of Fenton reaction, proteasome activity and iron-sulfur cluster biosynthesis in normoxic cells are higher than hypoxic cells (Figure S2). These observations match (1) decreased level of Fenton reactions under hypoxia condition due to the lack of oxygen and hydrogen peroxide, and (2) decreased levels of TCA cycle-related iron-sulfur cluster containing proteins. In addition, we have observed that the level of ferric iron reduction is largely suppressed in APEX1-KD cells, as the overrepresented ROS may deplete ferrous iron pool in the cell (Figure S2).

These observations demonstrated that the scFEA prediction can capture the major variations in iron ion metabolisms under different biochemical conditions. We have also conducted a robustness analysis by using TCGA data. Similar to our past validation of scFEA (50), we randomly shuffled the gene expression profile of each iron ion metabolic genes in a certain proportion of samples and evaluated the total loss with respect to the level of perturbation. We observed higher total losses when perturbing more samples, which further demonstrated that the iron ion metabolic gene expressions truly form certain dependency over the curated metabolic modules (Figure S3).

### Iron metabolism in human cancer

We have first applied scFEA on all the samples of nine cancer types and 11 subtypes (see Materials and Methods) vs. controls to predict the flux rates of the iron metabolism in Figure 1A. Key prediction results are summarized in **Figure 2**.

**Fig. 2.**
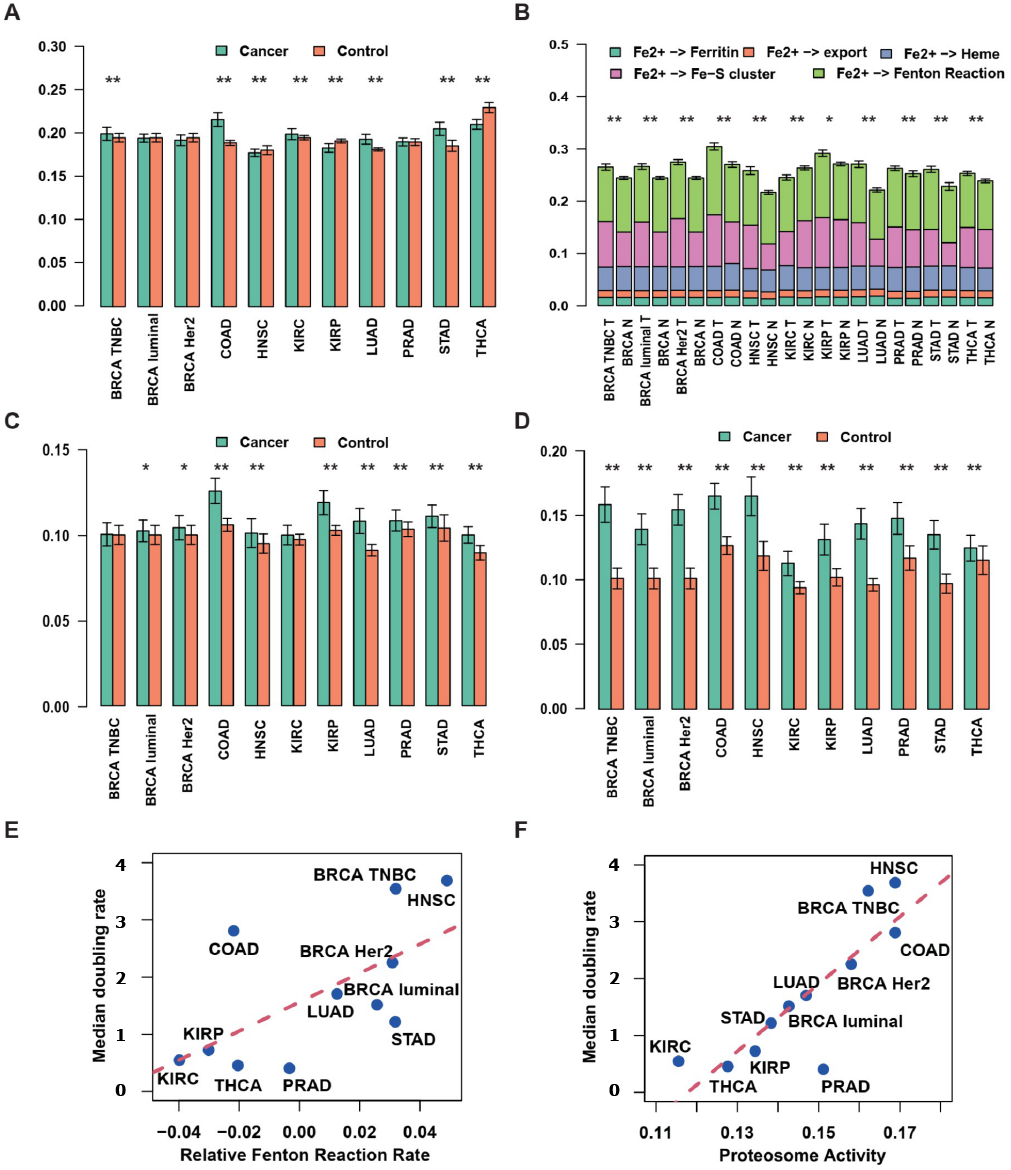
Predicted iron fluxes. The predicted fluxes are relative flux levels scaled by a hyperparameter. **A.** The (predicted) average iron import rate (y-axis) in cancer and adjacent control samples of each cancer type (x-axis). **B.** Predicted average cytosolic Fe2+ metabolic rates (y-axis) by ferritin synthesis, mitochondrial heme and Fe–S cluster synthesis, Fe^2+^ export, and Fenton reaction, in cancer and adjacent control samples of each cancer type (x-axis). **C.** Predicted average cytosolic Fenton reaction level (y-axis) in cancer and adjacent control samples of each cancer type (x-axis). **D.** Predicted averaged proteasome level (y-axis) in cancer and adjacent control samples of each cancer type (x-axis). In (A–D), * and ** suggest p-value < 0.1 and p-value < 0.05, respectively. **E.** Correlation between the difference of relative Fenton reaction rates in cancer and control tissues (x-axis) and cancer growth rate (y-axis). **F.** Correlation between the predicted proteasome activity level (x-axis) and cancer growth rate (y-axis).

Figure 2A summarizes the predicted uptake level of iron. Overall, five cancer (sub)types have elevated uptake level, namely BRCA_TNBC (*P* = 0.043), COAD (*P* = 2*E* – 16), KIRC (*P* = 0.02), LUAD (*P* = 1*E* – 10), and STAD (*P* = 1*E* – 7); three have reduced iron uptake level, namely HNSC (*P* = 0.02), KIRP (*P* = 1*E* – 5), and THCA (*P* = 3*E* – 15); and three have approximately the same level: BRCA_Luminal, BRCA_HER2, and PRAD. We have also predicted the levels of five exits for cytosolic Fe^2+^, namely ferritin synthesis, heme, Fe-S cluster, Fe^2+^ export, and Fenton Reaction, in cancer vs. controls, as shown in Figure 2B. Overall, we note that 10 out of the 11 cancer (sub)types have higher cytosolic Fe^2+^ exit level in cancer compared to their adjacent controls (*P* < 1*E* – 3, except for KIRP), which is mostly due to the increased Fe – S biosynthesis and Fenton reaction, as detailed in Figure 2B. Furthermore, all the 11 cancer subtypes have increased Fenton reaction rates compared to control tissues (Figure 2C), among which COAD (*P* = 6*E* – 18), HNSC (*P* = 0.02), KIRP (*P* = 9*E* – 12), LUAD (*P* = 8*E* – 21), PRAD (*P* = 0.004), STAD (*P* = 0.01), and THCA (*P* = 9*E* – 17) show significant changes while BRCA_TNBC, BRCA_Luminal, BRCA_HER2, and KIRC show slight but insignificant increases. On average, our prediction suggests that the Fenton reaction involves 39%-44% of the cytosolic Fe^2+^ utilization across the 11 cancer subtypes while more than 95% of the produced Fe^3+^ stored in ferritin. Statistics of all the iron ion metabolic modules in the TCGA cancer types are provided in Table S4.

We have also evaluated the level of the 20S proteasome which degrades particularly protein aggregates formed due to reaction with hydroxyl radicals (56) generated by Fenton reactions. Significantly increased proteasome level has been observed in all cancer types compared to controls (*P* < 1*E* – 5, Figure 2D), providing an independent support for that cancer tissues have higher levels of cytosolic Fenton reactions than controls, knowing that hydroxyl radicals can only be produced intracellularly by Fenton reactions. The cytosolic Fenton reaction level could be related to the cell growth rate (Figure 2E-F), which is discussed in detail below.

### RMs induced for proton production by alkalizing intracellular pH

We have previously hypothesized that RMs observed in cancer tissue cells of the same cancer type are predominantly induced by cytosolic Fenton reactions to neutralize the hydroxides produced by the reactions. The rationale for this hypothesis is a highly significant observation that two totally unrelated sets of reactions, namely cytosolic Fenton reactions and the total hydroxides produced by the observed RMs are strongly statistically correlated (41). In addition, multiple evidence strongly suggests that these two sets of reactions are causally linked and furthermore, Fenton reactions drive the induction of the RMs observed in each tissue sample but the other way around. Specifically, Fenton reaction is solely the result of increased innate immune responses, giving rise to increased H_2_O_2_ concentration and local iron overload and intracellular sequestration. In addition, none of the H^+^-producing RMs studied contribute to increased immunity or iron overload, based on our extensive literature review. Furthermore, OH^−^-producing Fenton reaction form a natural driver for the simultaneous induction of numerous reprogrammed metabolisms, which are distinct across different cancer types, to keep the intracellular pH stable. The other way around might require a biological program, which is orders of magnitude more complex than our current model. Based on these considerations, we have tested our hypothesis using our more reliable way for estimating the quantities of the involved reaction rates.

43 RMs are used here with their names and marker genes given in **Table 1**, including amino-acid biosynthesis and degradation, purine and pyrimidine biosynthesis, lipid and fatty acid biosynthesis and others. For each RM, its level is assessed using single-sample Gene Set Enrichment Analysis (ssGSEA) on individual samples (see Materials and Methods) (57).

**Table 1.**
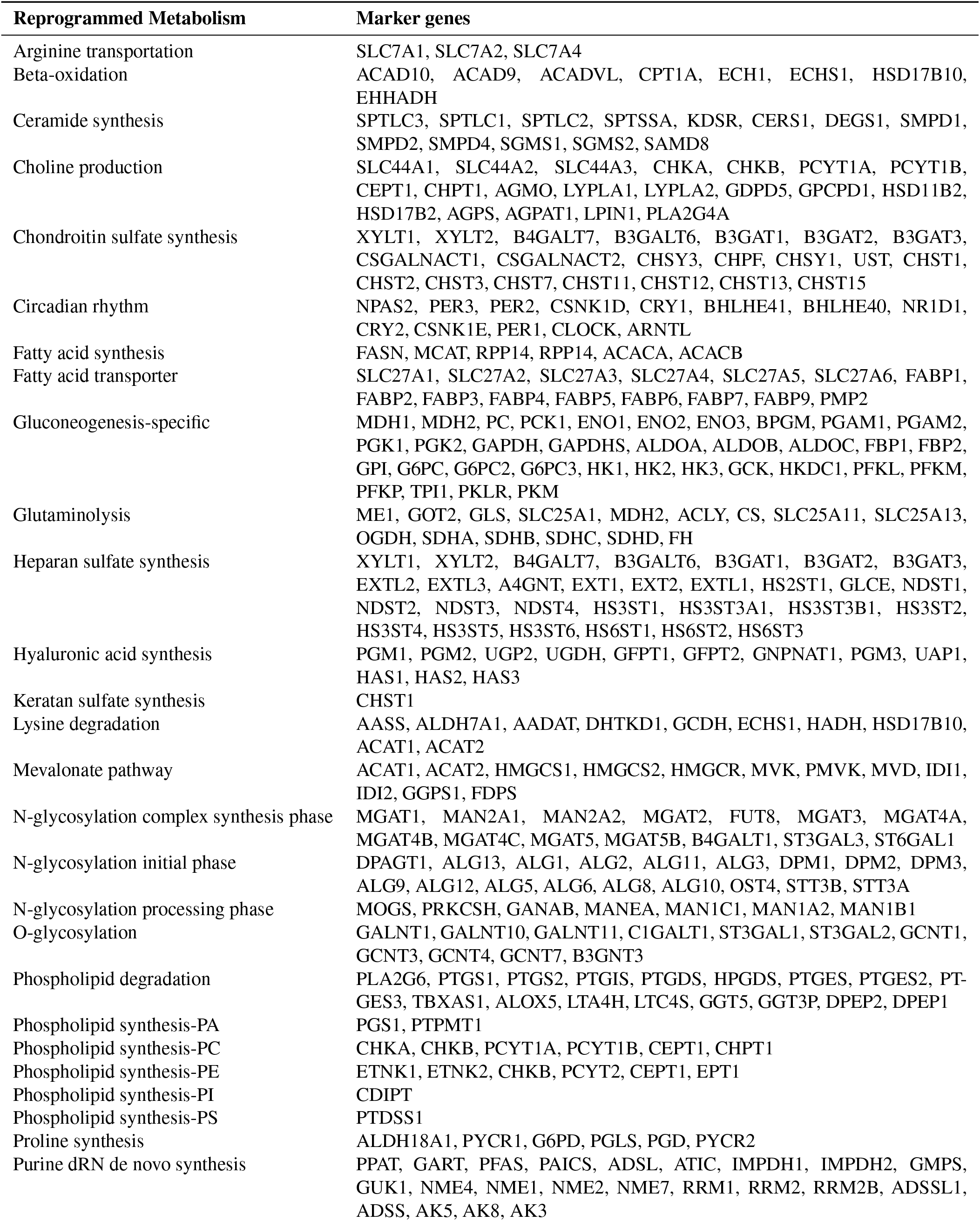
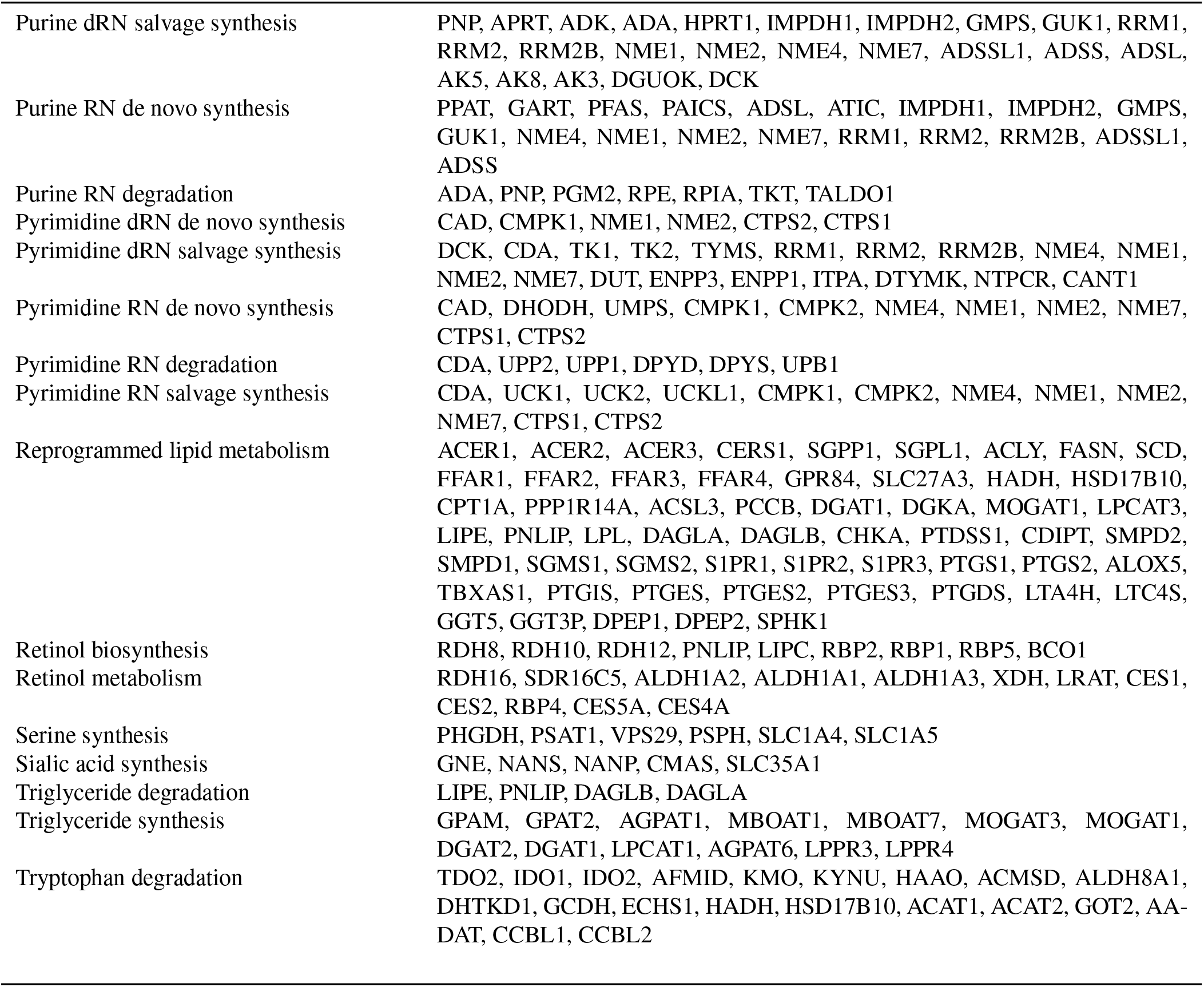
43 reprogramed metabolisms with names and marker genes

A linear regression of the predicted rate of the cytosolic Fenton reaction against the levels of the 43 RMs across all samples of the nine cancer types (and 11 subtypes) with the L1 penalty for variable selection (see Materials and Methods). **Table 2** shows the RMs, along with the numbers of H^+^ and CO_2_ produced, that are commonly and positively associated with Fenton reaction rates across all the samples, where the averaged contribution score and the rate of contribution represent the averaged regression parameter and the proportion of cancer types that the RMs were selected, respectively. Table S5 gives the selected RMs for each of the 11 cancer subtypes.

**Table 2.** Contribution scores of RMs that are positively correlated with Fenton reaction rates

| Reprogrammed metabolisms | Averaged contribution score | Rate of contribution | Net proton |
| --- | --- | --- | --- |
| Purine dRN salvage synthesis | 0.287 | 0.818 | +1 per purine |
| Proline synthesis | 0.23 | 1 | +1 CO<sub>2</sub> per proline |
| Tryptophan degradation | 0.112 | 0.818 | +1 per tryptophan |
| Pyrimidine RN salvage synthesis | 0.094 | 0.727 | +1 per pyrimidine |
| Pyrimidine dRN salvage synthesis | 0.091 | 0.636 | 0 or +1 per pyrimidine |
| Phospholipid synthesis-PE | 0.06 | 0.636 | +1 CO<sub>2</sub> per PE |
| Phospholipid synthesis-PA | 0.054 | 0.818 | +1 per PA |
| Phospholipid synthesis-PI | 0.053 | 0.636 | +1 per PI |
| Phospholipid synthesis-PS | 0.052 | 0.455 | +4 per PS |
| Phospholipid degradation | 0.047 | 0.727 | 0 or +1 per phospholipid |
| Mevalonate pathway | 0.041 | 0.545 | +1 CO<sub>2</sub> per farnesyl diphosphate |
| Gluconeogenesis-specific | 0.041 | 0.545 | +1 per pyruvate |
| Sialic acid synthesis | 0.041 | 0.636 | +2 per sialic acid |
| Fatty acid transporter | 0.033 | 0.909 | +1 per fatty acid |
| Beta-oxidation | 0.032 | 0.455 | +1 per fatty acid |

Note: $+n$ in column 4 denotes $n$ protons produced. PE, phosphatidylethanolamine; PA, phosphatidic acid; PI, phosphatidylinositol; PS, phosphatidylserine.

Positive associations of the following RMs were identified in at least 40% of cancer types: purine deoxyribonucleotide (dRN) salvage synthesis, proline synthesis, tryptophan degradation, pyrimidine ribonucleotide (RN) salvage synthesis, pyrimidine dRN salvage synthesis, phospholipid synthesis, phospholipid degradation, mevalonate pathway, gluconeogenesis-specific, sialic acid synthesis, fatty acid transporter, and beta-oxidation, hence possibly representing the most commonly selected RMs in all cancers.

The selected RMs together achieve higher than 0.8 *R*^2^ in explaining the Fenton reaction rate across all samples of the 11 cancer subtypes (**Figure 3**). Specifically, purine, pyrimidine and proline synthesis and tryptophan degradation have the strongest associations with the predicted Fenton reaction rate and are commonly induced in more than 80% cancer types. It is noteworthy that nucleotide biosynthesis represents the most powerful acidifier, knowing that *de novo* synthesis of a purine produces 8–9 net protons and that of pyrimidine produces 3–5 net protons per nucleotide. Proline synthesis is known to accelerate the glycolysis pathway (58) and also an effective producer of acids, as detailed below (41):

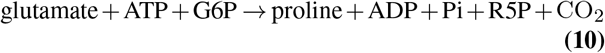

which produces one CO_2_ per proline synthesized. Cancer generally utilizes a truncated tryptophan degradation pathway, whose end-product is kynurenine or 3-hydroxyanthrranliate rather than the usual acetyl-CoA for the full degradation pathway. There could be two possible reasons for the employment of the truncated pathway, one being that this process produces net protons and the other being that both end-products promote cell survival under immune attacks (59).

**Fig. 3.**
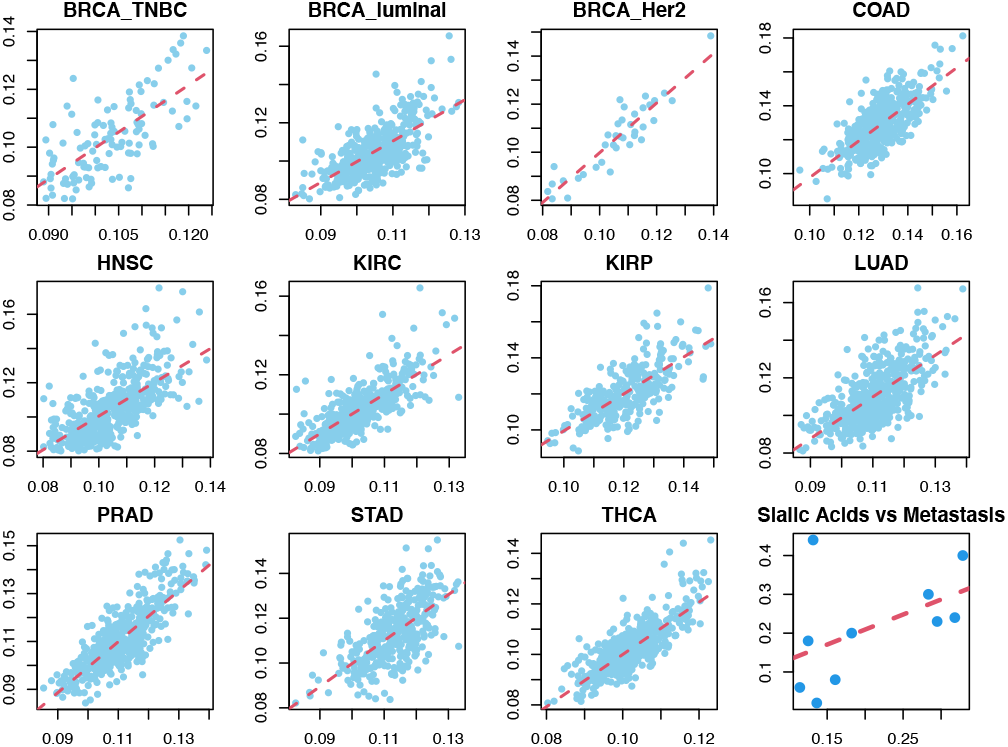
Fenton reaction rate vs H^+^ producing RMs. The first 11 panels each show a scatter plot with the predicted Fenton Reaction (y-axis) rate vs. the repression model prediction (x-axis) in 11 cancer (sub)types. The bottom right panel shows the known cancer metastasis rate (y-axis) vs the against sialic acid synthesis rate and sialic acid degradation gene *NEU1*’s expression.

### Linking cancer phenotypes to Fenton reaction levels and induced RMs

We aim to elucidate possible relationships between the phenotypes of a cancer and the RMs induced, knowing that cellular phenotypes are dictated by the metabolisms of the cell.

#### Cancer growth rate and cytosolic Fenton reaction level

For each of the 11 cancer subtypes, we have collected the average time needed for a tumor to double its volume, as detailed in **Table 3**. We have observed a strong positive correlation, Pearson Correlation Coefficient (PCC) = 0.635 (*P* = 0.036) (Figure 2E), between the increment in the relative Fenton reaction rates in cancer tissues vs. adjacent control tissues, defined as the proportion of Fenton reaction among the total flux of the five cytosolic Fe^2+^ outfluxes (Figure 2B), and the tumor-growth rate defined as 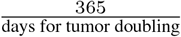. In addition, a stronger correlation was observed between the averaged 20S proteasome level and tumor growth rate, at PCC = 0.838 (*P* = 0.001) (Figure 2F). These provide strong evidence that the cytosolic Fenton reaction level plays a key role in dictating the rate of cancer growth.

**Table 3.** Average time needed to double the tumor size across 11 cancer subtypes

| Cancer type |  | Median doubling time (days) |
| --- | --- | --- |
| BRCA | TNBC | 103 |
|  | ER+ | 241 |
|  | HER2 | 162 |
| COAD |  | 10 |
| HNSC |  | 99 |
| KIRC |  | 667 |
| KIRP |  | 504 |
| LUAD |  | 214 |
| PRAD |  | 900 |
| STAD |  | 300 |
| THCA |  | 803 |

#### Cancer metastasis and sialic acid accumulation

Previous studies have suggested that the high-level of sialic-acid accumulation on cancer surface is associated with high metastasis rate. We have collected the metastasis rate of each cancer type under consideration and the synthesis of sialic acids. A positive correlation, PCC =0.55 (*P* = 0.09), between the combined predicted sialic acids synthesis and degradation rate and the metastasis rate of a cancer has been observed, hence providing strong evidence to the aforementioned speculation. In addition, we have also conducted a regression analysis to fit the cancer type-specific metastasis rate against the expressions of sialic acid synthesis and degradation gene *NEU1*, giving rise to the following relationship:

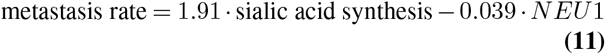

with p-values 0.071 and 0.076 for the two contributors, respectively. Hence the analysis suggests a positive association between metastasis rate and sialic acids synthesis and a negative association with sialic acid degradation, which together implies the rate of accumulation (Figure 3).

#### Local immune and stromal cell populations and Fenton reactions

For each cancer type, we have selected the 25% of the samples with the highest Fenton reaction levels, termed as samples with *high Fenton reactions*; and do the same on the 25% samples with the lowest Fenton reactions, termed as samples with *low Fenton reactions*. To study if the levels of cytosolic Fenton reactions may be associated with certain immune and stromal cell types, we have applied ICTD (identification of cell types and deconvolution), an in-house deconvolution method (60), to estimate the relative proportion of immune and stromal cells of different types in a cancer samples of the nine cancer types. Our previous analysis demonstrated ICTD could robustly identify and estimate the relative proportion of 21 immune and stromal cell types and proportions of the populations by each cell type in TCGA samples (see Materials and Methods).

In all the nine cancer types, the samples with high Fenton reactions tend to have fewer stromal cells, namely fibroblast cells, endothelial cells, muscle cells, adipocytes, and neural cells (**Figure 4** A). And such samples are negatively associated with the CD4+ T-cells and cytokine releasing neutrophils, all compared to the samples with low Fenton reactions. Furthermore, samples with high Fenton reactions have higher MHC class-I/II expressing cells, total T-cells, total B-cells, granulocytes and cytotoxic CD8+ T-cells (Figure 4A).

**Fig. 4.**
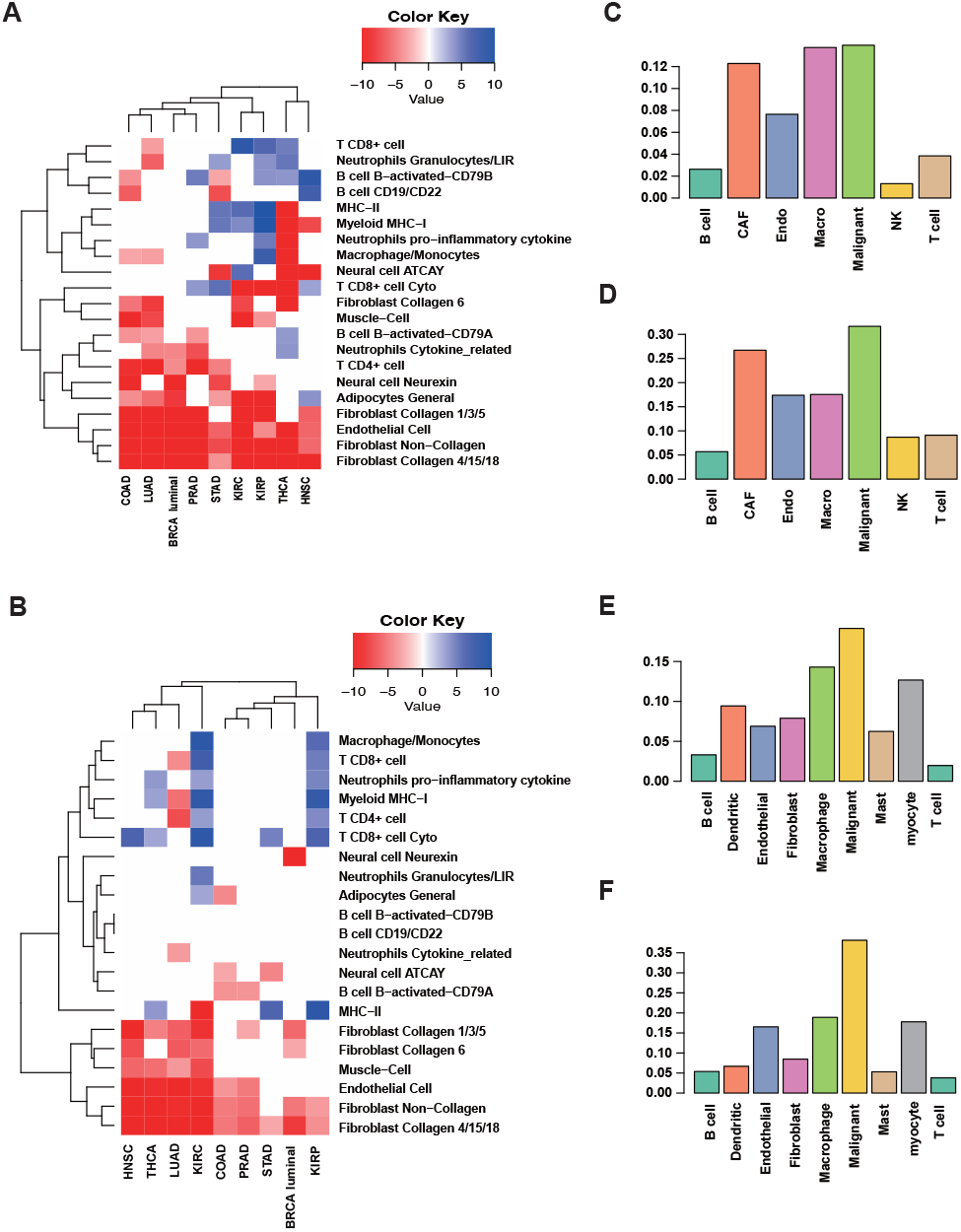
Variations of tumor microenvironments associated with Fenton reaction and OH^−^ level. **A.** Differences in immune and stromal cell populations between samples of high and low Fenton reaction rates. **B.** Differences in immune and stromal cell populations between samples of high and low OH-levels. **C–D.** Predicted Fenton reaction rates and proteasome levels in each cell type in the GSE72056 dataset. **E–F.** Predicted Fenton reaction rates and proteasome levels in each cell type in the GSE103322 dataset. The y-axis of **C–F** represents predicted relative flux rates.

We also compared the immune and stromal cell populations between the cancer samples with high and low OH^−^ levels (Figure 4B). We note that cancer samples with high OH^−^levels are negatively associated with stromal cell populations and positively associated with MHC class II antigen presenting cells, total T and B-cells, especially the cytotoxic CD8+ T-cell.

We have further conducted similar analyses on single-cell RNA-seq of human melanoma and head and neck cancer (HNSC). Our prediction suggests that in both cancer types, cancer cells have the highest Fenton reaction level (Figure 4C, E) and the proteasome level (Figure 4D, F) among all cell types, which confirms the above bulk data-based predictions using the TCGA samples. Other cell type specific iron metabolic fluxes of these two datasets were provided in Figures S4 and S5.

## Discussion

Acid-base homeostasis and its persistent disruption are known to play key roles in, possibly at the roots of, the development of a wide range of chronic illness ranging from type II diabetes, Alzheimer’s disease, Parkinson’s disease (9, 11–13) to cancer. Here we have provided strong evidence that such disruption of the intracellular pH, resulted from chronic inflammation and local iron accumulation, has a driving role in the induction of a range of RMs for cell survival. Based on our analysis results, each cancer (sub)type utilizes a unique combination of RMs at specific levels (Table S5), together serving as a stabilizer of the disrupted intracellular pH. Our modeling results strongly suggest that these induced RMs have given rise to the distinct phenotype of each cancer (sub)type. This, for the first time, provides a unified and effective framework for studying all the RMs and cancerous behaviors in a systematic manner.

While our analysis is largely correlation-based, it has strong causal implications since Fenton reactions are the results of chronic inflammation, and such reactions precede a majority of the metabolic reprogramming in diseases (manuscript in preparation). With this understanding, our framework can be considered as that Fenton reactions drives metabolic reprogramming, which determines the altered cellular behaviors. In this sense, we state that Fenton reactions may dictate a specific cancerous behavior. Note that we have focused on cytosolic Fenton reactions in this study while Fenton reactions in other subcellular locations, namely mitochondria, extracellular matrix and cell surface as we have previously suggested (28), may lead to some other cancerous behaviors, which we did not discuss here.

Our analyses have demonstrated that all the nine cancer types have selected *de novo* nucleotide biosynthesis as one of the top acidifiers to keep the intracellular pH stable. We have observed that most of the nine cancer types utilize biosynthesis and deployment of sialic acids and gangliosides as a key acidifier. In addition, lipid metabolism was another major acidifier in a few cancer types.

Our previous work has provided strong evidence that the rate of *de novo* biosynthesis of nucleotides may dictate the rate of cell proliferation (28), hence explaining why different cancer (sub)types may have distinct proliferation rates as shown in the Results section. In addition, the rate of sialic acid accumulation has strong implications to the rate of cancer metastasis (61), hence giving an explanation of why non-small cell lung cancer, kidney renal papillary cell carcinoma, head and neck cancer, and HER2+ breast cancers tend to have higher metastasis rates compared to others. Furthermore, different compositions of immune and stromal cell types in the cancerous tissues are associated with different levels of Fenton reactions. We anticipate that a variety of other phenotypes of a cancer could also be naturally explained using this framework, possibly once Fenton reactions in other subcellular locations being considered, such as the levels of resistance to different drugs, the possible secondary locations of metastasizing cancers, and the possibilities of development of cachexia.

It should be noted that this framework not only provides a capability for explaining why a cancer (sub)type has specific phenotypes in terms of the induced RMs, but also enables studies of the possible relationships among different phenotypic characteristics of a cancer such as growth vs. metastatic rates. For example, we have learned from the Results section that the relationship between the rates of cancer cell proliferation and metastasis could be strongly correlated with nucleotide biosynthesis and sialic acid synthesis, respectively, which tend to serve as the top and the dominating acidifiers in cancer. Hence for a given level of hydroxide production in a cancer, a relatively higher level of nucleotide biosynthesis may imply a lower level for the sialic acid synthesis since they together are utilized as the key acidifier. We expect that similar arguments can be made about the relationships among other top acidifiers for a given cancer type.

### Fenton reaction and ferroptosis

Both local iron overload and Fenton reactions have been long and widely observed across numerous cancer types (28). A natural question is: do cancer tissue cells tend to have ferroptosis? We have noted that the key enzymes, ACSL4, LPCAT3 and ALOX15, for converting polyunsaturated fatty acids (PUFAs) to PUFA-OOH, the main source of lethal lipid peroxides, whose production leads to ferroptosis, tend to be downregulated for at least two out of the three enzymes in seven out of nine cancer types except for HNSC and STAD, both of which have two enzymes upregulated, as detailed in Table S6. Further analyses have revealed that the levels of the cytosolic Fenton reactions have negative Spearman correlations with the marker genes for the cellular response to hydroperoxides except for LUAD, as detailed in Table S7.

We have also analyzed the differential expressions in tissues samples at different stages I–IV based on the clinical data retrieved from TCGA. At least one of the three key markers for ferroptosis, *ACSL4*, *LPCAT3*, and *ALOX15*, is downregulated in all cancer types and stages except for COAD and STAD. On average, 44 out of the 66 Fenton reaction marker genes are upregulated in all cancers and stages. These differentially expressed genes also display a trend: the genes become more up- or downregulated in advanced stages than in early stages. Detail statistics of the differential expression of Fenton reaction and ferroptosis related genes are given in Table S8.

These indicate that cytosolic Fenton reactions in general do not contribute to but possibly prevent the production of lethal lipid peroxides, which leads to ferroptosis. Knowing that cytosolic Fenton reactions generally take place in either labile iron pool or in some iron-containing proteins like heme in cancer and hydroxyl radical generally travels no more 1 nano meter (62), we speculate that the hydroxyl radicals produced by Fenton reactions may not reach lipids, say in the membrane in cancers, and may even take away some Fe2+ from taking part in lipid peroxidation as our statistics suggest.

### Perspective

A key realization from the study here is that the links from chronic inflammation and iron overload to pH-related stress and then to induced RMs and further to phenotypical features of each cancer type may represent the backbone of the development of a cancer while other changes such as genomic mutations and epigenomic alterations may predominantly serve as facilitators for realization of this evolutionary process as some authors have suggested (63), including our own (41). Compared to signaling and regulatory processes, metabolic events are considerably more stable as shown by common RMs shared by multiple cancer types. An important implication is that the issue of “drug resistance” could be potentially avoided by focusing on metabolisms rather than signaling/regulatory processes since the issue of “drug resistance” essentially reflects the redundance (or robustness) nature of the signaling or regulatory processes in human cells and organs, hence suggesting a possibly new direction of enzyme-centric cancer treatment, i.e., to inhibit key enzymes that acidify the cancer intracellular space and hence kill the cells.

It is noteworthy that this study examines the acid-base homeostasis and RMs from a perspective of chemical balances. We did not touch on issues related to the signaling and regulatory processes that connect disrupted homeostasis and induction of certain RMs nor touch on roles played by genomic mutations as well as epigenomic activities in these inductions and their downstream activities. These could represent as future research directions to provide further mechanistic information about the functional role played by signaling and regulatory molecules in the induction of acidifying metabolisms. A related issue is to elucidation of the possible reasons for why certain metabolisms are induced reprogrammed in some cancer types but not in other types, hence possibly leading to deepened understanding about specific cancer types and specific cancerous behaviors.

## Materials and methods

### Data used in this study

#### TCGA transcriptomic data

TCGA RNA-seq v2 FPKM data of the nine cancer types (11 subtypes) were retrieved from the Genomic Data Commons (GDC) data portal using TCGAbiolinks (64). **Table 4** lists the names of the cancer types along with the information of the numbers of cancer and control samples. FPKM values were converted to Transcripts per Million (TPM) values as the latter is more stable across samples. Clinical data were obtained in XML format from GDC and parsed with an in-house script. GENCODE gene annotations used by the GDC data processing pipeline was downloaded directly from the GDC reference files webpage.

**Table 4.** Tumor and normal sample sizes in each cancer type

| Abbreviation | Cancer type | Tumor sample count | Normal sample count |
| --- | --- | --- | --- |
| BRCA | Breast invasive carcinoma | 1091 | 113 |
| COAD | Colon adenocarcinoma | 456 | 41 |
| HNSC | Head and neck squamous cell carcinoma | 500 | 44 |
| KIRC | Kidney renal clear cell carcinoma | 530 | 72 |
| KIRP | Kidney renal papillary cell carcinoma | 288 | 32 |
| LUAD | Lung adenocarcinoma | 513 | 59 |
| PRAD | Prostate adenocarcinoma | 495 | 52 |
| STAD | Stomach adenocarcinoma | 375 | 32 |
| THCA | Thyroid carcinoma | 502 | 58 |

#### Single cell RNA-seq data

We have collected two scRNA-seq datasets from the GEO database, namely:

GSE72056: This dataset is collected on human melanoma tissues. The original paper provided cell classification and annotations including B-cells, cancer-associated fibroblast cells, endothelial cells, macrophage cells, malignant cells, NK cells, T-cells, and unknown cells (65).
GSE103322: This dataset is collected on head and neck cancer tissues. The original paper provided cell classification and annotations including B-cells, dendritic cells, endothelial cells, fibroblast cells, macrophage cells, malignant cells, mast cells, myocyte cells, and T-cells (66). Notably, as indicated by the original work, malignant cells have high intertumoral heterogeneity. Basic processing was conducted by using Seurat (version 3) (67) with default parameters to filter out cells with high expressions of mitochondrial genes. The cell type label and sample information provided in the original work were directly utilized.

### Software and statistical methods

#### ssGSEA

We applied the ssGSEA2.0 R package to estimate the levels of the selected RMs on individual samples (57). The ES score computed by ssGSEA was utilized to represent the level of each RM. Gene sets of the RMs were collected and annotated in our previous work (50).

#### scFEA

We have applied our scFEA method (50) on the TCGA and two scRNA-seq data against the iron metabolic map.

#### Regression analysis of Fenton reaction rate vs. RM levels

We have conducted a linear regression to fit the Fenton reaction rate against the RM levels across all samples of each cancer type. GLMnet R package was utilized for the regression analysis (68). An L1 penalty was utilized for variable selection. The hyperparameter lambda was determined through cross validation. The RMs positively associated with the Fenton reaction rate in at least 40% of the cancer types under study were summarized.

#### Samples with high and low Fenton reactions and OH^−^ levels

We have extracted the top and bottom 25%samples in terms of their predicted Fenton reaction level in each cancer type as cancer type specific high and low Fenton reaction samples. Similarly, we have done that in terms of the OH^−^ levels.

#### Deconvolution analysis

We have utilized our in-house deconvolution method, ICTD (identification of cell types and deconvolution) to estimate the relative proportions among 21 immune and stromal cell types in each TCGA sample (60).

#### Statistical test of differential analysis

We have utilized Mann Whitney test for all differential analysis, including differential gene expression analysis and the difference of predicted flux.

## Data availability

The TCGA data used in this study can be downloaded from the Genomic Data Commons (https://portal.gdc.cancer.gov/). Two scRNA-Seq data sets were collected from the GEO database with accession numbers GSE72056 and GSE103322.

## Code availability

The code for performing the analyses in this study can be found at https://github.com/changwn and https://github.com/y1zhou.

## CRediT author statement

**Yi Zhou**: Methodology, Software, Formal analysis, Visualization, Writing – Original Draft, Writing – Review & Editing. **Wennan Chang**: Methodology, Software, Formal analysis, Visualization, Writing – Original Draft. **Xiaoyu Lu**: Software, Data Curation. **Jin Wang**: Methodology, Validation, Writing – Review & Editing. **Chi Zhang**: Methodology, Validation, Supervision, Writing – Original Draft, Writing – Review & Editing. **Ying Xu**: Conceptualization, Validation, Supervision, Writing – Original Draft, Writing – Review & Editing.

## ACKNOWLEDGEMENTS

This work is supported by the National Science Foundation of USA (Grant No. 2047631) and partially by Georgia Research Alliance.

## Supplementary material

**Figure S1 Fenton reaction levels in cancer and non-cancerous chronic disease tissues.** The y-axis shows the log2(fold change) of 20S proteasome genes in disease versus control tissues. RNA-Seq data of chronic disease tissues were retrieved from GSE123661, GSE163416, GSE159008, and GSE84346. Differential expression analysis was performed using DESeq2.

**Figure S2 Predicted flux of nine selected modules under four conditions.** The conditions shown on the x-axis are normoxic APEX1 knockdown (si), normoxic control (sc), hypoxic APEX1 knockdown (si_h), and hypoxic control (sc_h).

**Figure S3 Total loss of scFEA computed with different ratios of samples perturbed.** The x-axis shows the ratio of samples perturbed. The baseline loss is shown as a red dashed line. Here the perturbation was conducted by first randomly selecting a certain proportion of samples and then randomly shuffling each row of the samples.

**Figure S4 Predicted flux of six selected modules in human melanoma.** The x-axis shows the eight cell types from the GSE72056 human melanoma scRNA-seq data set.

**Figure S5 Predicted flux of six selected modules in human head and neck cancer.** The x-axis shows the nine cell types from the GSE103322 human head and neck cancer scRNA-seq data set.

**Table S1 Average expression levels of *SLC9A1* across all samples in control and four stages of each of nine cancer types.**

**Table S2 Average expression levels of *CA4* and *CA7* in control and each stage in nine cancer types.**

**Table S3 Genes used to estimate the flux of the 15 metabolic modules.**

**Table S4 P values-of the difference of each iron ion metabolic module between each (sub)cancer type vs. normal control sample.**

**Table S5 Selected reprogrammed metabolisms for each of the 11 cancer subtypes.**

**Table S6 Average expression levels of three key enzyme genes involved ferroptosis.**

**Table S7 Correlation coefficients between Fenton reaction level and hydroperoxide marker genes.**

**Table S8 Log2 fold-change of Fenton reaction and ferroptosis related genes.**

