## Supplementary figures and images for "Acid-Base Homeostasis and Implications to the Phenotypic Behaviors of Cancer"

### Figure S1

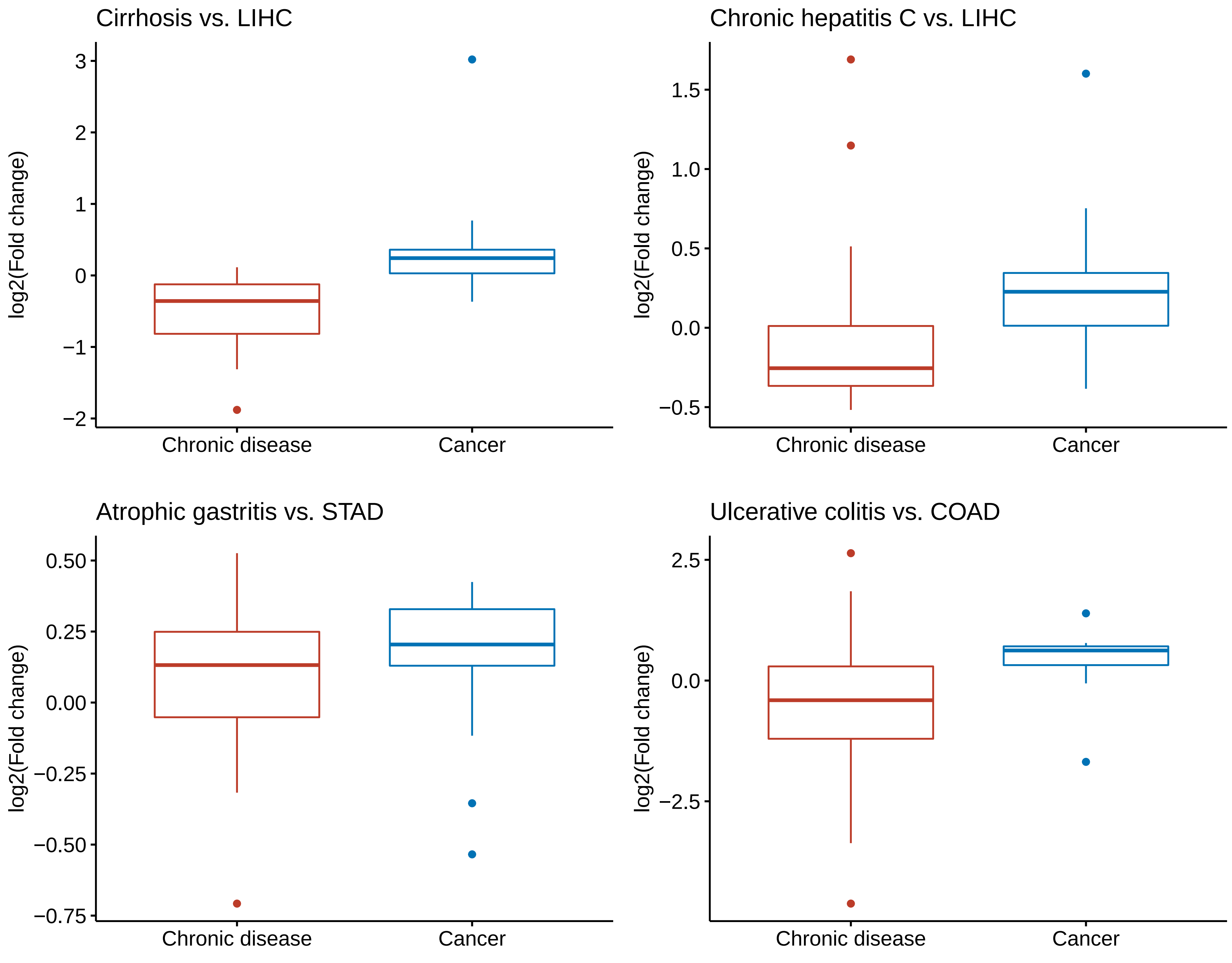

### Figure S2

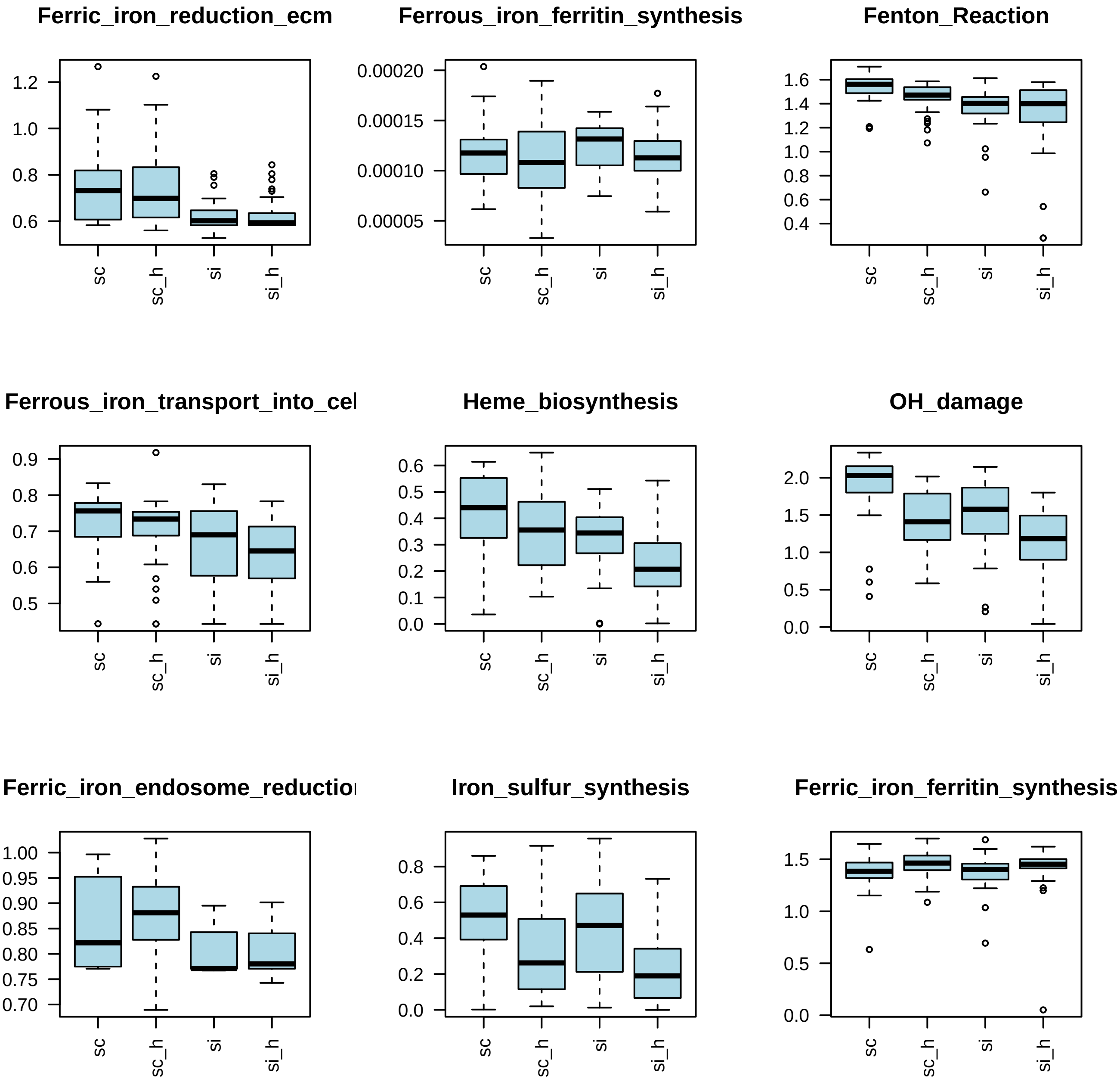

### Figure S3

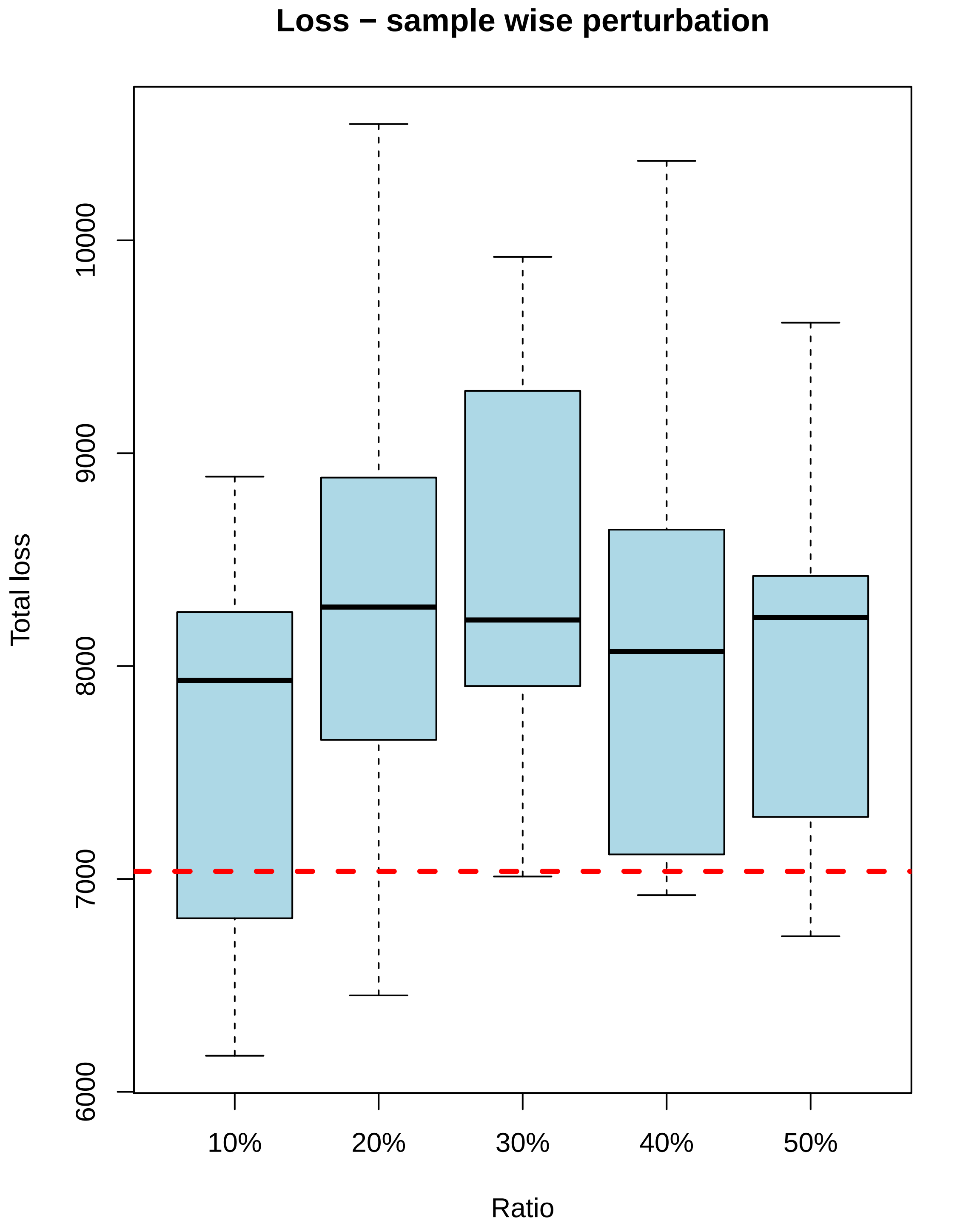

### Figure S4

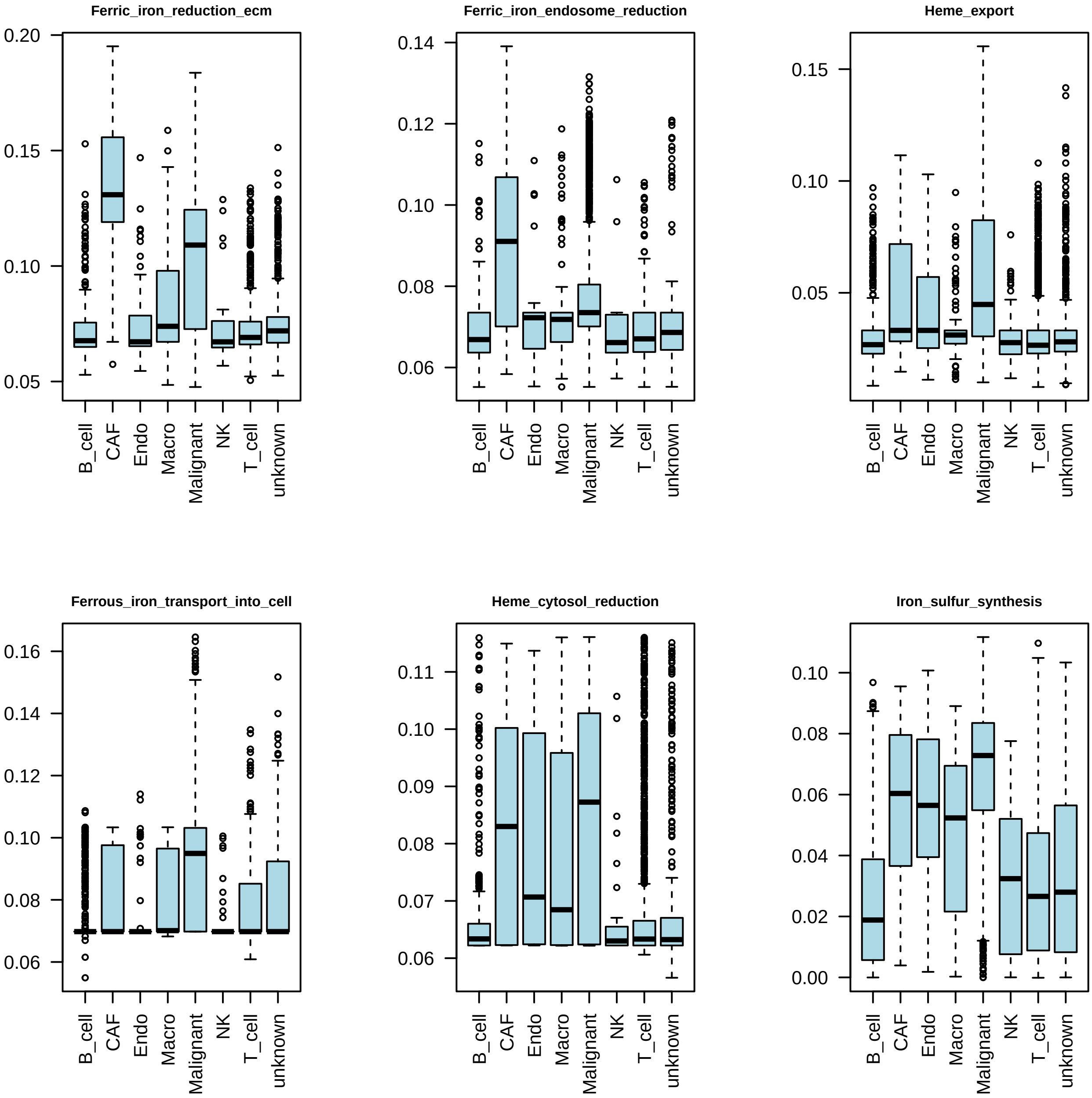

### Figure S5

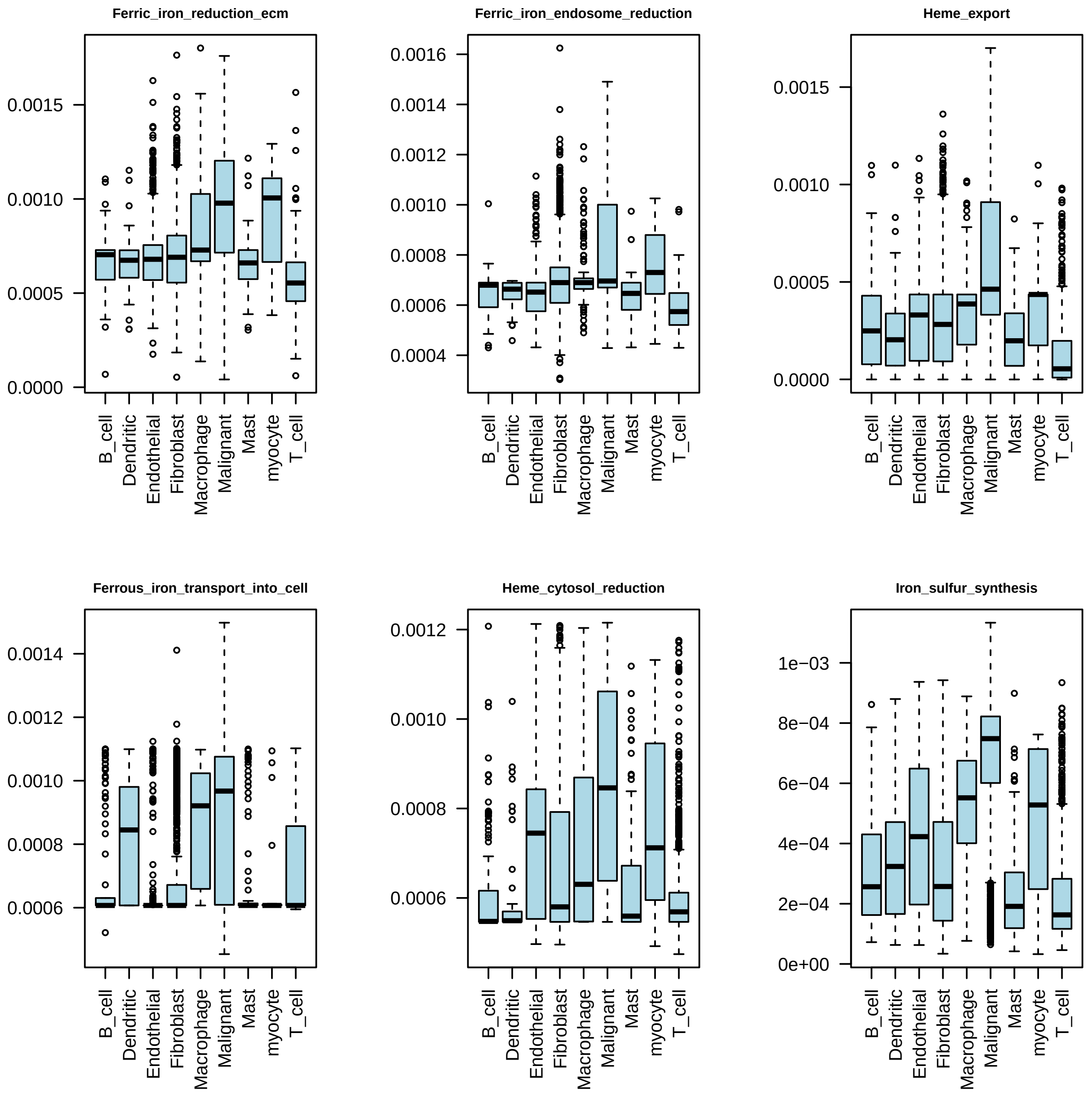
